# A Recombinant Antibody Against Human DRP1 Serine 616 Phosphorylation Enables Detection of BRAF^V600E^-Associated Mitochondrial Division in Cancer

**DOI:** 10.64898/2026.01.16.699897

**Authors:** Shanon T. Nizard, Yiyang Chen, Madhavika N. Serasinghe, Ruben Fernandez-Rodriguez, Kamrin D. Shultz, Jesminara Khatun, Anthony Mendoza, Juan F. Henao-Martinez, Jesse D. Gelles, Stella G. Bayiokos, J. Andrew Duty, Shane Meehan, Mihaela Skobe, Jerry Edward Chipuk

## Abstract

Mitochondria are dynamic organelles that continuously undergo balanced cycles of fusion and division to meet cellular demands. Mitochondrial division is mediated by Dynamin-Related Protein 1 (DRP1), a cytosolic large GTPase whose phosphorylation at serine 616 (DRP1-S616Ⓟ) promotes its translocation to the outer mitochondrial membrane and organelle division. Dysregulated, mitochondrial division disrupts cellular homeostasis and contributes to disease pathogenesis, including cancer. Our prior work demonstrated that the oncogene-induced mitogen-activated protein kinase (MAPK) pathway constitutively phosphorylates DRP1 at serine 616 (DRP1-S616Ⓟ), which is essential to cellular transformation and correlates with oncogene status in patient tissues. Similarly, DRP1-S616Ⓟ is subject to pharmacologic control by targeted therapies against oncogenic MAPK signaling. Building upon this foundation, we developed a human recombinant monoclonal antibody with high specificity for DRP1-S616Ⓟ, referred to as 3G11. Using diverse biochemical platforms, we demonstrate the robust utility of 3G11 to detect DRP1-S616Ⓟ in melanoma cell extracts and isolated organelles. Immunofluorescence revealed that pharmacologic inhibition of oncogenic MAPK signaling reduces DRP1-S616Ⓟ levels which correlates with mitochondrial hyperfusion; while immunohistochemistry showed that elevated DRP1-S616Ⓟ expression in human tissues correlates with BRAF^V600E^ disease. Together, these findings establish 3G11 as a specific, versatile, renewable, and cost-effective tool for studying mitochondrial division, with strong potential for clinical applications.

## 1. Introduction

Mitochondria are double membrane organelles that undergo regulated cycles of fusion and division to maintain cellular and organismal homeostasis [3]. Dysregulation of these dynamic processes contributes to various pathological conditions, such as inflammation, metabolic disorders, neurodegeneration, and cancer [4,5]. Mitochondrial dynamics is primarily coordinated by large GTPases: Mitofusin 1 (MFN1) and Mitofusin 2 (MFN2) mediate outer mitochondrial membrane (OMM) fusion, while Optic Atrophy 1 (OPA1) controls inner mitochondrial membrane (IMM) fusion [6]. Mitochondrial division is mediated by Dynamin Related Protein 1 (DRP1), a monomeric cytosolic protein that translocates to the OMM upon activation, binds to adaptor proteins (*e.g.,* Mitochondrial Fragmentation Factor, MFF), and oligomerizes to mediate GTP-dependent scission of both the OMM and IMM [2].

Mitochondrial fusion ensures the proper exchange of lipids, metabolites, mtDNA, and proteins between mitochondria which is essential for maintaining organelle function [1]. Mitochondrial division is critical for the distribution of mitochondria to daughter cells during mitosis, subcellular localization of mitochondria to meet local metabolic demands, and facilitating mitochondrial quality control [2]. The regulation of DRP1-dependent mitochondrial division is a central regulator of disease biology across multiple organ systems. When DRP1 is hyperactivated or suppressed, it directly impacts metabolism, apoptosis, inflammation, and cell fate. As such, DRP1 is subject to a variety of post-translational mechanisms (*e.g.,* phosphorylation) which regulate DRP1 activation, membrane translocation, and oligomer stabilization [7].

The Mitogen Activated Protein Kinase (MAPK) pathway is a master regulator of proliferation, differentiation, and survival that directly controls DRP1 function [8]. Previous studies suggest a mechanistic connection between oncogene-induced MAPK (*e.g.,* RAS^Q61R^, BRAF^V600E^) signaling and mitochondrial dynamics. Specifically, ERK1/2 (p42/p44) phosphorylates DRP1 at serine 616 (DRP1-S616Ⓟ), thus activating DRP1 translocation to the OMM and mitochondrial division [11]. Oncogene-induced chronic mitochondrial division leads to impaired mitochondrial function, characterized by reduced oxidative phosphorylation and an increased reliance on alternative metabolic pathways. Literature suggests that DRP1-dependent mitochondrial division is essential to cancer mechanisms as silencing DRP1 inhibits transformation [11]. Furthermore, pharmacological inhibition of MAPK signaling in RAS and BRAF mutant human cancer models eliminates DRP1-S616Ⓟ, resulting in hyperfused mitochondria, re-activation of mitochondrial function, and sensitization to cell death [11]. DRP1-S616Ⓟ is observed in tumors with oncogenic MAPK mutations and is suggested to support the development and growth of breast, colon, glioblastoma, lung, pancreatic, and skin cancers [12–17].

In the majority of cutaneous melanomas, the BRAF^V600E^ mutation occurs early in pathogenesis but is insufficient to induce malignancy and requires additional events to initiate disease [19]. One downstream event subsequent to BRAF^V600E^ induced transformation is sustained DRP1-S616Ⓟ suggesting chronic mitochondrial division may be a robust biomarker for BRAF^V600E^ melanoma and progression. Indeed, DRP1-S616Ⓟ status dichotomized with BRAF^V600E^ and was detected in ∼80% of nevi that subsequently progressed to melanoma, where it persisted throughout the melanomagenesis process [20]. This persistence highlights the potential for DRP1-S616Ⓟ as a reliable indicator for disease initiation and a promising tool for improving our understanding of melanoma biology. However, the majority of antibodies raised against DRP1-S616Ⓟ are traditional monoclonals with undefined variable heavy/light regions (VH/L), restricted by commercial availability and cost, and the early generation hybridomas are subject to genetic drift potentially impacting their specificity. To circumvent these constraints, we developed a recombinant monoclonal DRP1-S616Ⓟ antibody, referred to as 3G11. We validated the 3G11 antibody using a range of techniques, including immunoblotting, immunofluorescence, and immunohistochemistry in melanoma models and patient tissues. These data support the specificity of 3G11 for DRP1-S616Ⓟ and highlighted its potential utility in detecting oncogene-induced chronic mitochondrial division in melanoma and diverse disease models and patient samples.

## 2. Materials and Methods

### 2.1. Reagents

Opti-MEM Reduced Serum Media (Gibco, Cat. No. 31985-070) was supplemented with 5% FBS, 1% penicillin/streptomycin, 0.01 µg/mL heparin (Sigma-Aldrich, Cat. No. H3393), 0.01 µg/µL human FGF-basic (Gibco, Cat. No. PHG0026), 0.05 mg/mL dibutyryl cyclic-AMP sodium salt (Sigma-Aldrich, Cat. No. DO627), and 0.1 mM 3-isobutyl-1-methylxanthine (Sigma-Aldrich, Cat. No. I5789). 2% Tumor Media was prepared with L-15 (Leibovitz) medium (Sigma Aldrich, Cat. No. L486) and supplemented with 2.5 µg/mL insulin solution (Sigma Aldrich, Cat. No. I0516), 0.187 mg/mL calcium chloride dihydrate (Sigma Aldrich, Cat. No. C7902), 1.5 mg/mL sodium bicarbonate (Santa Cruz Biotechnology, Cat. No. 203271A), 17.53 g/L MCDB 153 powder (Sigma Aldrich, Cat. No. L1518-500ML), 1% penicillin/streptomycin, and 2% FBS. Following preparation, 2% Tumor Media was filtered using a 0.2 micron sterile Nalgene filter unit with polyethersulfone (ThermoFisher-Scientific, Cat. No. 567-0020). Dulbecco’s Modification of Eagle’s Medium (DMEM) (Corning, Cat. No. 10-017-CV) was supplemented with 3% FBS, 1% penicillin/streptomycin, and 1% L-glutamine. The drugs used in this study are GSK1120212 (Selleck Chemicals, Cat. No. S2673) and PLX4032 (Selleck Chemicals, Cat. No. S1267). The antibodies utilized for western blot are total DRP1 (Cell Signaling Technology, Cat. No. 8570), DRP1-S616Ⓟ (Cell Signaling Technology, Cat. No. 3455S), total ERK (Cell Signaling Technology, Cat. No. 137F5), ERK1/2Ⓟ (Abcam, Cat. No. AB278538), β-Actin (Abcam, Cat. No. ab8227), TOMM20 (Santa Cruz Biotechnology, Cat. No. SC-17764), MFN1 (Santa Cruz Biotechnology, Cat. No. SC-166644), MFN2 (Santa Cruz Biotechnology, Cat. No. SC-515647), and OPA1 (Invitrogen, Cat. No. MA5-32786). For microscopy, cells were stained for HSP60 (Cell Signaling Technology, Cat. No. 12165) or TOMM20 (Santa Cruz Biotechnology, Cat. No. SC-17764), Alexa Fluor® 594-conjugated goat anti-mouse IgG secondary antibody (ThermoFisher-Scientific, Cat. No. A11032), and Alexa Fluor® 488-conjugated goat anti-rabbit IgG secondary antibody (ThermoFisher-Scientific, Cat. No. A11034). Cells were co-stained with Hoechst 33342 (ThermoFisher-Scientific, Cat. No. H3570) to visualize nuclei. For immunohistochemistry, tissues were stained with recombinant 3G11 and BRAF^V600E^ (Cell Signaling Technology, Cat. No. 2900S).

### 2.2. Cell Culture

The YUPEET human-derived primary melanoma cell line was cultured in Opti-MEM Reduced Serum Media. The WM35, WM853, WM902, WM983, and WM1152 human-derived primary melanoma lines were cultured in 2% Tumor Media. A375 and SKMEL28 human-derived non-primary (*i.e.,* lymph node and secondary metastasis derived, respectively) melanoma lines were purchased from ATCC and cultured in DMEM media. All cell lines were cultured at 37°C in a humidified 5% CO_2_ incubator, passaged every 3-4 days, and regularly tested for mycoplasma contamination.

### 2.3. Immunizations and Hybridoma Generation

For hybridoma generation, five 8-week-old BALB/c female mice were primed subcutaneously (s.c.) with 80 µg (40 µg in each rear flank) of a keyhole limpet hemocyanin (KLH) conjugated DRP1 peptide spanning amino acid residues 607–620, in which Serine 616 was phosphorylated (KLH-Cys-SKPIPIMPA{pSer616}PQKG) (GenScript USA Inc., Piscataway, NJ) in the presence of complete Freund’s adjuvant (CFA). Four weeks after priming, a total of two more boosts commenced in the same manner in the presence of incomplete Freund’s adjuvant. After immunizations, blood was collected from the submandibular vein before each boost to monitor titer of sera antibodies by enzyme-linked fluorescent immunosorbent assay (ELISA).

For the ELISA, plates were coated with 10 mg of non-KLH conjugated phosphorylated peptide in 1× PBS followed by blocking with 1% BSA/PBS and washing (3×) using a BioTek 405 microplate washer (Agilent Technologies). Sera were diluted in 1% BSA/PBS and added to the coated well. After 3× washes, goat anti-mouse IgG-HRP (1:4000 in 1% BSA/PBS) (Jackson ImmunoResearch Labs, West Grove, PA; Cat. No. 115-035-003) was added and positive wells were detected with ABTS substrate solution (Roche; Cat. No. 11684302001). All incubation steps were performed at 25°C for 1 hour. Unimmunized sera were also added as negative control. The mouse with the highest titer at the 1:10,000 dilution that was also >2× the negative control was selected for hybridoma fusion and received 2 final boosts of 150 µg BSA-conjugated phosphorylated-DRP1 peptide s.c. (50 µg in 3 sites) over 2 days. A final bleeding (“fusion sera”) and terminal splenectomy were performed 3 days after the last final boost. The spleen was processed to a single-cell suspension and fused in the presence of ClonaCell HY-PEG (Stemcell Technologies, Vancouver, BC; Cat. No. 03806) with Sp2/0 hetero-myeloma fusion partners. Individual hybridoma B cell clones were grown under selection on soft agar containing hypoxanthine-aminopterin-thymidine (HAT), and expanded colonies were picked ten days later into 96 well tissue culture plates using a robotic ClonaCell Easy Pick instrument (Hamilton/Stem Cell Technology) in ClonaCell HY-Medium E (Stemcell Technologies; Cat. No. 03805). Clones were expanded, and the supernatant was used to screen for DRP1 detections. Animals were housed in an Association for the Assessment and Accreditation of Laboratory Animal Care approved facility. The animals were provided with enrichment and cared for by qualified animal technicians. All animal studies were approved by the Icahn School of Medicine at Mount Sinai’s Institutional Animal Care and Use Committee (IACUC), IACUC #IPROTO202200000097 (Nov. 11, 2025 approved), and the animal studies strictly adhered to ARRIVE guidelines.

### 2.4. Dot Blot Analysis

Indicated peptides were solubilized in 1×PBS, spotted onto nitrocellulose membranes, and the membranes were dried. Membranes were rehydrated in 1× PBST and blocked in 4% nonfat dry milk in 1× PBST. Membranes were incubated with either the indicated anti-serum or commercially available DRP1 and DRP1-S616Ⓟ antibodies diluted in blocking buffer (1:1000, 4% milk in 1× PBST) at 4°C for 14 to 16 hours. After incubation, blots were washed with 1× PBST and visualized using enhanced chemiluminescence detection (Millipore, Cat. No. WBLUF0100).

### 2.5. Cloning and Expression of the 3G11 Antibody

Sequencing of the 3G11 antibody was performed using SMARTer 5’ RACE technology (Takara Bio, USA, Cat. No. 634858) adapted for immunoglobulins to amplify the variable genes from the heavy and kappa chains from the 3G11 hybridoma. Briefly, RNA was extracted from the hybridoma using a RNeasy Mini Kit (Qiagen, Cat. No. 74104), followed by first-strand cDNA synthesis using constant gene-specific 3’ primers (GSP1) based on mouse IgG/mouse kappa constant isotypes and incubation with the SMARTer II A Oligonucleotide and SMARTscribe reverse transcriptase (Takara).

GSP1 Primers (5’–3’):

mG1-AGAGGTCAGACTGCAGGACA, mG2a-CTTGTCCACTTTGGTGCTGC,

mG2b-GACAGTCACTGAGCTGCTCA, mG2b-GACAGTCACTGAGCTGCTCA, and mcK-CCAACTGTTCAGGACGCCAT.

PCR amplification of the first-strand cDNA product was then performed using SeqAmp DNA Polymerase (Takara, Cat. No. 638504) with a nested 3’ primer (GSP2 Primer) to the constant genes and a 5’ universal primer (kit provided) based on universal primer sites added to the 5’ end during cDNA generation.

GSP2 Primers (5’–3’):

mG1-CCCAGGGTCACCATGGAGTT, mG2a-GGTCACTGGCTCAGGGAAAT, mG2b-CTTGACCAGGCATCCCAGAG, mG3-GACAGGGCTCCATAGTTCCATT, and mCk-CTGAGGCACCTCCAGATGTTAAC.

Purified PCR products were submitted for Sanger sequencing using 3’ constant gene primers (GeneWiz, South Plainfield, NJ). Sequence results were BLASTed against the IMGT mouse databank of germline genes using V-Quest (http://imgt.org/). Complete 3G11 heavy and kappa chain variable region cDNAs were then synthesized with an Il-2 signal sequence and cloned into a pcDNA-3.4 mammalian expression vector in-frame on the 3’ end to a vector-encoded mouse IgG2a constant region (heavy chain) or a mouse kappa constant region (light chain) (GenScript USA Inc., Piscataway, NJ). Cloned vectors representing full heavy and kappa chains were then transfected in a 1:1 ratio into HEK Expi293 suspension cells using ExpiFectamine transfection reagent following manufacturer instructions (ThermoFisher, Cat. No. A14524). Supernatant was collected 5 days after initial transfection, filtered (0.45 micron), and subjected to quantitation/verification by bio-layer interferometry (BLI) and purification.

### 2.6. Quantitation and Verification of 3G11 Supernatant

For BLI analysis, 3G11 supernatant was quantitated and verified on an Octet Red96 (ForteBio, Sartorius). Optically capable biosensors conjugated with Protein A were dipped into supernatants containing the 3G11 mAb. Supernatants were measured undiluted and diluted 1:10 in conditioned media and compared to an isotype standard (mG2a), diluted in conditioned media in a 1:2 dilution series ranging from 100 µg/mL to 1.56 µg/mL. Standard curves were analyzed in the ForteBio Data Analysis Software v.10 (ForteBio) using a 5 parameter logistics (5PL) dose response curve fitting model on the initial binding slopes. MAb concentrations are then calculated from the standard curve. Diluted samples were compared to undiluted samples after application of the respective dilution factors. Total concentrations were averaged together from the diluted and undiluted sample.

### 2.7. 3G11 Antibody Purification

The supernatant was purified on an ÄKTA pure FPLC systems using a Protein-A affinity columns (HiTrap-1 mL, GE/Cytiva, #17-0404-01) with UV monitored extraction. A single UV peak fraction was collected representative of the antibody, dialyzed against PBS and quantitated by both bicinchoninic assay (BCA) (Pierce Biotechnology, Cat. No. 23225) and absorption at OD_280nm_.

### 2.8. In Vitro Kinase Assay and Analysis

Recombinant, GST-tagged, full-length, human DRP1 protein (10 ng/µL) was incubated with recombinant ERK1 kinase (2 ng/µL; Sigma, Cat. No. 14-439-M) in kinase buffer (50 mM Tris-HCl pH 7.4, 30 mM NaCl, 15 mM MgCl₂, 200 µg/mL BSA, 2 mM DTT, and 1 mM ATP) at 37°C for 30 minutes. The reaction was terminated by the addition of 4× SDS loading buffer (0.2 M Tris-HCl, 0.4 M 2-mercaptoethanol, 277 mM SDS, 6 mM Bromophenol Blue, 4.3 M Glycerol). DRP1 (100 ng/lane) was subjected to SDS-PAGE, followed by standard western blotting. The nitrocellulose membrane was blocked in 5% dried nonfat milk in 1× TBST and incubated with primary antibodies (1:1000; 5% milk in 1× TBST) at 4°C for 14 to 16 hours. The membrane was then washed 3 times with 1× TBST, incubated with the secondary antibody (1:2500; 5% milk in 1× TBST) at 25°C for 1 hour, washed 3 times with 1× TBST, and detected using enhanced chemiluminescence (Millipore, Cat. No. WBLUF0100).

### 2.9. Whole Cell Lysate Isolation and Analysis

One 10-cm dish of cells at 80% confluency was used per condition. Cells were treated with DMSO, GSK1120212 (50 nM), or PLX4032 (1 µM) for 6 hours. Cells were trypsinized, pelleted by centrifugation at 800 *× g* for 10 minutes, washed with pre-chilled 1× PBS, and pelleted again using the same parameters. The cell pellets were lysed by resuspending in 1× RIPA buffer supplemented with protease inhibitors (ThermoFisher-Scientific, HALT Tablet, Cat. No. 87786) and phosphatase inhibitors (ApexBio, Cat. No. k1015b). The suspension was incubated on ice for 10 minutes and then centrifuged at 21,000 *× g* at 4°C for 10 minutes. Protein concentrations were determined using a BCA assay. Cell lysates were adjusted with 1× RIPA buffer to achieve the same protein concentration across samples. 4× SDS loading buffer(0.2 M Tris-HCl, 0.4 M 2-mercaptoethanol, 277 mM SDS, 6 mM Bromophenol Blue, 4.3 M Glycerol) was added to each sample and diluted to a final 1× concentration, Samples were then boiled at 95°C for 5 minutes. 75 µg of cell lysate per lane was subjected to SDS-PAGE followed by transferring to a nitrocellulose membrane. The nitrocellulose membrane was blocked in 5% milk/1× TBST and then incubated with primary antibodies (1:1000, 5% milk in 1× TBST) at 4°C for 14–16 hours. The membrane was then incubated with the secondary antibody (1:2500, 5% milk in 1× TBST) at 25°C for 1 hour before enhanced chemiluminescence detection (Millipore, Cat. No. WBLUF0100).

### 2.10. Heavy Membrane Isolation and Analysis

At least two 15-cm dishes of cells at 90–95% confluency were used per condition for heavy membrane (*i.e.,* mitochondrial fractions) isolations. Cells were treated with DMSO, GSK1120212 (50 nM), or PLX4032 (1 µM) for 6 hours. The cells were harvested by trypsinization and pelleted by centrifugation at 800 *× g* at 4°C for 10 minutes. The cell pellets were washed with 1 mL of pre-chilled 1× PBS, pelleted, washed with 1 mL of pre-chilled trehalose isolation buffer (TIB) [300 mM Trehalose, 10 mM HEPES-KOH (pH 7), 10 mM KCl, 1 mM EDTA, 1 mM EGTA], and pelleted by centrifugation at 800 *× g* for 5 minutes. The cell pellets were resuspended with TIB (supplemented with 1× protease inhibitors, ThermoFisher-Scientific, HALT Tablet, Cat. No. 87786) at a 1:1 volume and incubated on ice for 30 minutes to allow for cell swelling. The suspension was then transferred into a pre-chilled 2 mL Potter-Elvehjem dounce and homogenized with fifty strokes to reach 70– 80% cell lysis. The homogenate was then transferred to a 1.5 mL Eppendorf tube and centrifuged at 1,000 *× g* at 4 °C for 5 minutes. The supernatant was collected and centrifuged again under the same conditions to ensure that no unlysed cells or nuclei were present. The resulting supernatants were centrifuged at 10,000 *× g* at 4°C for 10 minutes, and the pellets were collected as the heavy membrane fraction. Sample yield was determined using the Pierce BCA Assay Kit, cell lysates were adjusted with 1× RIPA buffer to achieve the same protein concentration across all samples, and 4× SDS loading buffer was added. 50 µg of cell lysate was loaded per lane, and analyzed using SDS-PAGE and western blotting. See [21] for a more detailed protocol on mitochondria isolations.

### 2.11. Immunofluorescence (IF)

A375 and SKMEL28 cells were seeded on 1.5-mm coverslips (Electron Microscopy Science, Cat. No. 72230-01) and allowed to adhere for 24 hours. The next day, cells were treated with DMSO or GSK1120212 (50 nM) for 16 hours. After treatment, cells were washed 3 times with 1× PBS and fixed in 4% formaldehyde for 15 minutes. The cells were then washed another 3 times in 1× PBS and then permeabilized using 0.3% Triton X-100 at 25°C for 12 minutes. Cells were washed again 3 times with 1× PBS and subsequently incubated in the blocking buffer (50% 1× PBS, 4% BSA, and 10% goat serum) at 25°C for 2 hours. Coverslips were incubated with primary antibodies (3G11, 1:3200; TOMM20, 1:400; HSP60:, 1:400) diluted in the blocking buffer at 4°C for 14–16 hours. Secondary antibodies Alexa Fluor® 594 (1:250) and Alexa Fluor® 488 (1:250) were diluted in the blocking buffer and incubated at 25°C for 1 hour. Samples were treated with Hoechst 33342 (1:5000), mounted with ProLong Glass (ThermoFisher-Scientific, Cat. No. P36980) and cured at 4°C overnight.

Images were captured at the Mount Sinai Tisch Cancer Center Microscopy and Advanced Bioimaging Shared Resource. Images were acquired with LAS X software (Leica Application Suite X 4.3.0.24308) on Leica Stellaris 8 (Leica Microsystems GmbH, Wetzlar, Germany) confocal microscope. Imaging was performed using a 63×/1.3 HC PlanApo oil immersion lens (Leica Microsystems GmbH, Wetzlar, Germany), the frame size was set to 2048×2048 pixels (X/Y), and images were acquired at 8 bits. The acquisition was performed at frame scan mode, using 1 frame averaging and 2 line accumulation. Fluorophores were excited with a 440 nm laser derived from an 80 MHz pulsed White Light Laser (Leica Microsystems GmbH, Wetzlar, Germany). The fluorophore emission was collected with Hybrid Detectors (HyDS, Leica Microsystems GmbH, Wetzlar, Germany) with a spectral window of 445–650 nm. The laser power and gain were set to provide the best signal-to-noise ratio for each chromophore while utilizing the full dynamic range of the detector and avoiding detector saturation.

Image analysis and mitochondrial quantification were performed using ImageJ’s plugins MitochondrialAnalyzer-V2.3.1 and DeconvolutionLab2_2.0.0. All images were pre-processed manually, which included masking the image to focus on a single cell (if there are multiple cells in the frame), performing a background subtraction, deconvolution (if not done through microscopy software while imaging), and image thresholding. Background subtraction was performed with a 4-pixel rolling-ball radius to minimize the background noise while still maintaining the integrity of the mitochondrial structure. Deconvolution was performed through the DeconvolutionLab2 plugin and was optimized for these images in respect to resolution, morphology structure, image size, and overall image quality. A Gaussian PSF with a Richardson-Lucy algorithm (15 iterations) was used to develop the deconvolved image. Once deconvolved, all images were auto-thresholded using Moments from ImageJ. Moments creates a binary based on the pixel intensity in the original image histogram and is a very consistent thresholding option. Mitochondrial Analyzer2D Analysis provides per-mitochondrion and summary data for each cell, using the binary image to identify total count, branch count, branch length, and additional metrics. For the purposes of elongation and fragmentation, the area to perimeter and perimeter to area ratios were calculated to better illustrate the level of fragmentation between groups. A power test revealed that 21 cells were necessary to establish statistical significance at a 5% significance level, with a *t*-test used to evaluate *p*-values.

### 2.12. Immunohistochemistry (IHC)

De-identified formalin-fixed paraffin-embedded (FFPE) human skin samples biopsy sections (5 micron thickness) were obtained from the Mount Sinai Tisch Cancer Center Biorepository and Pathology Shared Resource. A Leica BOND RX automated system for immunohistochemistry was utilized to stain tissue samples using a Bond Polymer Refine Red Detection Kit (Leica Biosystems, Cat. No. DS9390). Primary antibodies (were diluted in BOND Primary Antibody Diluent (Leica Biosystems, Cat. No. AR9352) as indicated: 3G11 (1:1000) and BRAF^V600E^ (1:150). Bond Epitope Retrieval (ER) Solution 2 (Leica Biosystems, Cat. No. AR9640) was used for 20 minutes; the slides were then mounted with DAKO Glycergel mounting medium (DAKO, Cat. No. C0563) and imaged. Slides were scored by a binary measure (0 or 1) which represents the absence/presence of stain, and a subset of samples were confirmed by pathologists within the Mount Sinai Division of Dermatopathology. Scoring was analyzed via a Fisher’s Exact Test and a Chi-Squared Test with Yates correction. Statistics were performed with GraphPad Prism software. The Program for the Protection of Human Subjects office determined that the above study is exempt from human research (HS#13-00606) as defined by DHHS regulations (45CFR46.101(b)(4)).

## 3. Results

### 3.1. Antibody Development Workflow and Screening to Identify 3G11 Anti-Sera

To develop a recombinant monoclonal antibody that specifically detects DRP1-S616Ⓟ, we designed an immunization strategy using a keyhole limpet hemocyanin (KLH)-conjugated peptide corresponding to human DRP1 amnio acid residues 607-620, with serine 616 phosphorylated. This phospho-peptide served as the immunogen for mouse immunizations, and subsequent hybridomas were generated by fusing splenic B cells from immunized mice with Sp2/0 hetero-myeloma cells [22]. Once the hybridomas were screened, the optimal VH/L regions were sequenced, cloned, and purified; the workflow is summarized **(Figure 1A)**. The immunogen was determined based on the secondary structure of DRP1, and this region is highlighted in an AlphaFold predicted structure of full-length human DRP1 **(Figure 1B)**. To identify anti-sera capable of specifically recognizing the serine 616 phosphorylated epitope, we performed an ELISA assay comparing interactions between the DRP1-S616 and DRP1-S616Ⓟ peptides **(Figure 1C)**. Anti-sera that exhibited strong reactivity were further evaluated using dot blot analysis to confirm selectivity for the phosphorylated DRP1 sequence **(Figures S1A–C)**. Based on the specificity for the DRP1-S616Ⓟ peptide by ELISA and dot blot, we selected five mice: 2G9, 3G11, 5E1, 10C8, and 10D10 for hybridoma generation and further validation.

**Figure 1.**
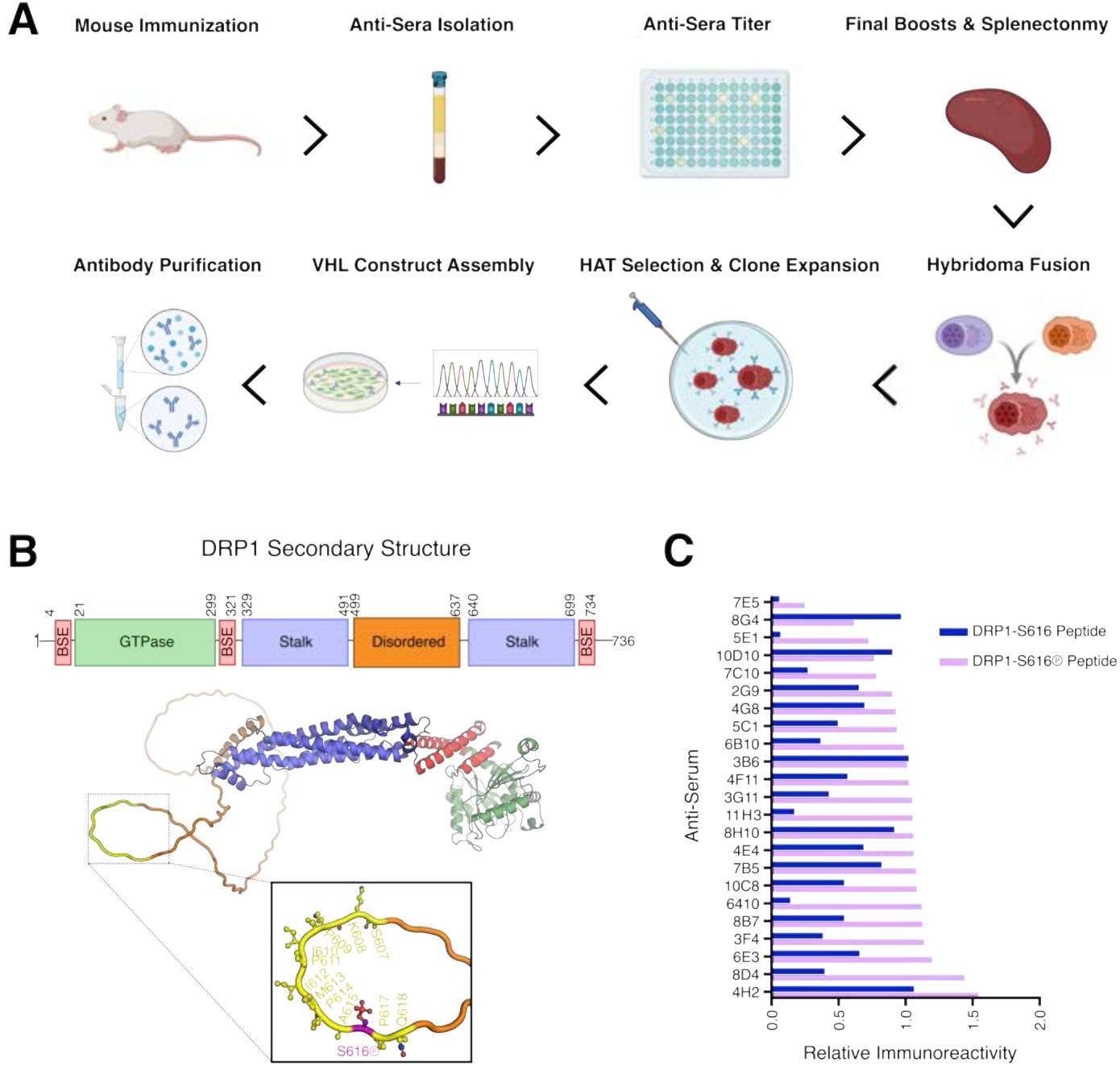
Antibody development workflow, DRP1 domains, and anti-sera screening. **(A)** Schematic representation of the workflow to generate the human, monoclonal, recombinant 3G11 antibody, including: immunization, anti-sera screening, hybridoma development, and identification. HAT and VH/L are Hypoxanthine/Aminopterin/Thymidine and Variable Heavy/Light chain, respectively. **(B)** Predicted three-dimensional structure of human DRP1 (AlphaFold V2, AF-O00429-v4), rendered using PyMOL (Schrodinger LLC.), with structural domains color coded based on domain definitions (Rochon et al., 2024). The epitope recognized by the 3G11 antibody is highlighted in yellow, and DRP1-S616Ⓟ is shown in purple. **(C)** ELISA results comparing the indicated anti-sera for reactivity to the DRP1 serine 616 non-phosphorylated and phosphorylated peptides.

We next determined which of these hybridomas distinguished between unphosphorylated and phosphorylated full-length, recombinant, human DRP1. First, we performed an *in vitro* kinase assay to specifically phosphorylate DRP1 at serine 616 using recombinant ERK1, as described [11,12]. DRP1-S616 and DRP1-S616Ⓟ were subjected to SDS-PAGE and western blot, and the 5 hybridomas were analyzed for specific binding **(Figure 2A)**.

**Figure 2.**
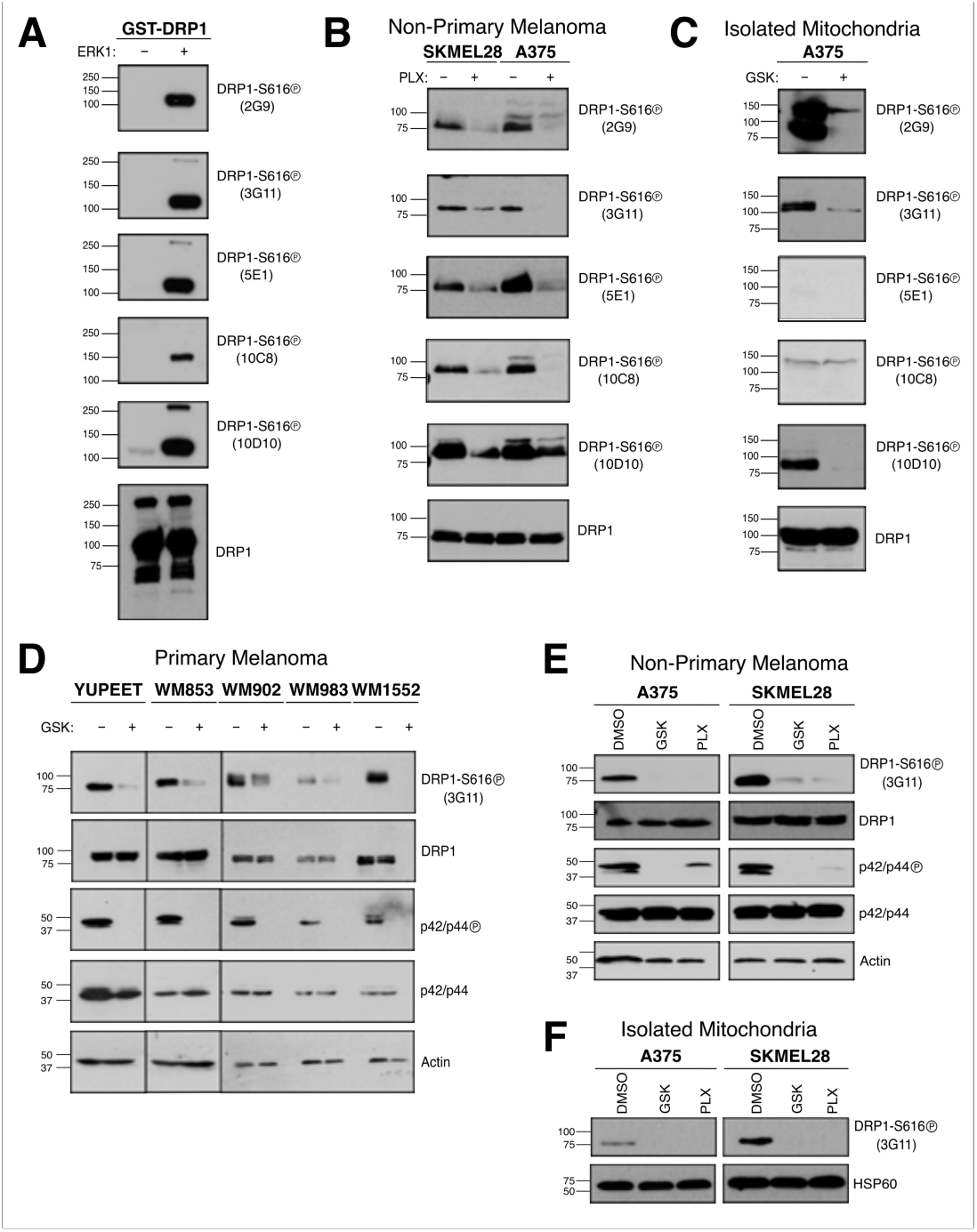
Hybridoma screenings and recombinant 3G11 antibody evaluation. **(A)** Hybridomas generated from the top five anti-sera were evaluated for specificity using western blot analysis of full-length recombinant GST-DRP1 ± ERK1 treatment. 100 ng of recombinant GST-DRP1 was loaded per lane. Total DRP1 was evaluated for equal protein loading. **(B–C)** The hybridomas were screened using western blot analysis of whole cell lysates (100 µg/lane; *B*) or heavy membrane fractions (25 µg/lane; *C*) isolated from SKMEL28 and A375 cells treated with GSK1120212 (50 nM; GSK), PLX4032 (1 µM; PLX), or DMSO for 6 hours. Total DRP1 was evaluated for equal protein loading. **(D)** Primary melanoma lines were treated with GSK1120212 (50 nM) or PLX4032 (1 µM) for 6 hours. Whole cell lysates were western blotted for indicated proteins. Recombinant 3G11 was evaluated for DRP1-S616Ⓟ detection. Actin was probed for equal protein loading. **(E)** A375 and SKMEL28 were treated with GSK1120212 (50 nM) or PLX4032 (1 µM) for 6 hours and evaluated as in *D*. Recombinant 3G11 was evaluated for DRP1-S616Ⓟ detection. Actin was probed for equal protein loading. **(F)** Heavy membrane fractions from the same treatments in *D* were western blotted for the indicated proteins. Recombinant 3G11 was evaluated for DRP1-S616Ⓟ. HSP60 was probed for equal protein loading. Molecular weight standards for all western blots are indicated in kilodaltons (kDa). Data are representative of 3 independent experiments.

Next, we assessed the ability of these hybridomas to detect endogenous DRP1-S616Ⓟ in whole cell lysates isolated from BRAF^V600E^ positive human malignant melanoma cell lines (*i.e.,* A375 & SKMEL28) **(Figure 2B)**. To functionally validate the clones’ specificities, we also treated these cells with PLX4032 (or DMSO vehicle), a targeted therapy against BRAF^V600E^, which inhibits downstream endogenous ERK1/2 from phosphorylating DRP1 at serine 616 [11]. Western blot analysis was performed to compare DRP1-S616Ⓟ levels in DMSO versus PLX4032-treated cells. Among the five hybridomas, 3G11 demonstrated the strongest and most-specific detection of DRP1-S616Ⓟ in DMSO-treated cells, with a potent reduction in DRP1-S616Ⓟ following PLX4032 exposure, indicating that 3G11 effectively detects endogenous DRP1-S616Ⓟ **(Figure 2B)**.

As DRP1-S616Ⓟ accumulates at the OMM [7], we subsequently evaluated which hybridoma could detect activated DRP1-S616Ⓟ localized to endogenous mitochondria. A375 cells were treated with GSK1120212 (or DMSO), a MEK inhibitor that suppresses ERK1/2, prior to isolating the mitochondrial fraction. SDS-PAGE and western blot analysis of isolated mitochondrial proteins revealed that the 3G11 anti-serum potently detected the presence of DRP1-S616Ⓟ in DMSO treated samples with minimal background; furthermore, there was an appreciable decrease in DRP1-S616Ⓟ detection following GSK1120212 treatment **(Figure 2C)**.

### 3.2. The Recombinant 3G11 Antibody Specifically Detects DRP1-S616 Phosphorylation

Through our screening process, we identified 3G11 as the optimal clone for detecting DRP1-S616Ⓟ based on its peptide specificity in ELISA and dot blot studies, and western blot analyses using whole cell lysates and isolated mitochondrial fractions (Figures 1 and 2). Therefore, we sequenced and cloned the 3G11 VH/L regions, and generated a purified recombinant monoclonal version of the 3G11 antibody. Using the purified recombinant 3G11 antibody, we proceeded to examine its ability to detect DRP1-S616Ⓟ in several melanoma models before and after oncogenic MAPK pathway inhibition.

To broaden our approach, we expanded our analyses to include a panel of primary melanoma cell lines harboring the BRAF^V600E^ mutation (*i.e.,* YUPEET, WM853, WM902, WM983, & WM1552) treated with GSK1120212 (or DMSO). SDS-PAGE and western blot analyses of whole cell lysates from these cell models revealed that ERK1/2 inhibition led to a significant reduction in 3G11 detection of DRP1-S616Ⓟ levels compared to DMSO-treated cells **(Figure 2D)**. Inhibition of MAPK signaling was also confirmed by a corresponding reduction in p42/p44 (ERK1/2) phosphorylation in all cell lines to ensure GSK1120212 treatments were effective. Together, these data indicate a selective effect of oncogenic MAPK pathway inhibition on DRP1-S616 phosphorylation.

Next, to further validate recombinant 3G11’s ability to detect DRP1-S616Ⓟ across different melanoma models, we extended our analysis to the non-primary BRAF^V600E^ melanoma cell lines, A375 and SKMEL28. As in figure 2B, we treated the cell lines with either GSK1120212 or PLX4032 and performed western blot analysis of whole cell lysates. In both cell lines, DRP1-S616Ⓟ levels decreased upon treatment with either inhibitor, while total DRP1 expression remained unchanged confirming that recombinant 3G11 specifically detects DRP1-S616Ⓟ **(Figure 2E)**.

We further evaluated whether recombinant 3G11 detected DRP1-S616Ⓟ in isolated mitochondrial fractions. Mitochondria were isolated from A375 and SKMEL28 cells treated with either GSK1120212 or PLX4032, and SDS-PAGE/western blot analysis was performed. In both cell lines, DRP1-S616Ⓟ levels decreased in the mitochondrial fractions following oncogenic MAPK inhibition, while total mitochondrial DRP1 remained unchanged **(Figure 2F)**. These results confirm that recombinant 3G11 specifically detects the presence of DRP1-S616Ⓟ and can be used to monitor its regulation in primary and nonprimary melanoma models using whole cell lysates and isolated mitochondria.

### 3.3. Recombinant 3G11 Detects Oncogenic MAPK-Regulated DRP1-S616 Phosphorylation and Mitochondrial Co-localization

To evaluate the utility of recombinant 3G11 in immunofluorescence (IF) based applications, we performed immunostaining using the BRAF^V600E^ non-primary melanoma cells (*n.b.,* A375 and SKMEL28 display more organized mitochondrial networks compared to the primary melanoma lines, which eases imaging analyses). SKMEL28 cells were treated with DMSO or GSK1120212 (50 nM, 16 hours), fixed, and immuno-stained using recombinant 3G11 to detect DRP1-S616Ⓟ, TOMM20 (OMM marker), and Hoechst 33342 (nuclear stain). First, we confirmed that GSK1120212 treatment inhibited mitochondrial division and resulted in elongated mitochondrial networks. Both standard 2D images and 3D projections of GSK1120212 treated SKMEL28 cells revealed a loss in mitochondrial division and elongated networks **(Figures 3A–B)**. This effect was corroborated with the quantification of the perimeter to area ratio **(Figure 3C)**. Next, we assessed the effects of oncogenic MAPK inhibition on DRP1-S616Ⓟ signal and its co-localization with mitochondria. In DMSO treated cells, DRP1-S616Ⓟ was prominently detected and exhibited colocalization with mitochondria **(Figure 3D)**. After GSK1120212 treatment, DRP1-S616Ⓟ expression was reduced, with a corresponding decrease in mitochondrial co-localization, as quantified using Mander’s coefficient **(Figures 3D–E)**. Similar results were obtained using the A375 melanoma cell line, where GSK1120212 treatment led to mitochondrial elongation and decreased DRP1-S616Ⓟ signal **(Figures 3F–G)**. Together, these findings validate the use of recombinant 3G11 in high-resolution imaging of DRP1-S616Ⓟ in melanoma cells and demonstrate that 3G11 reliably reports changes in DRP1-S616Ⓟ localization and mitochondrial dynamics in response to oncogenic MAPK inhibition.

**Figure 3.**
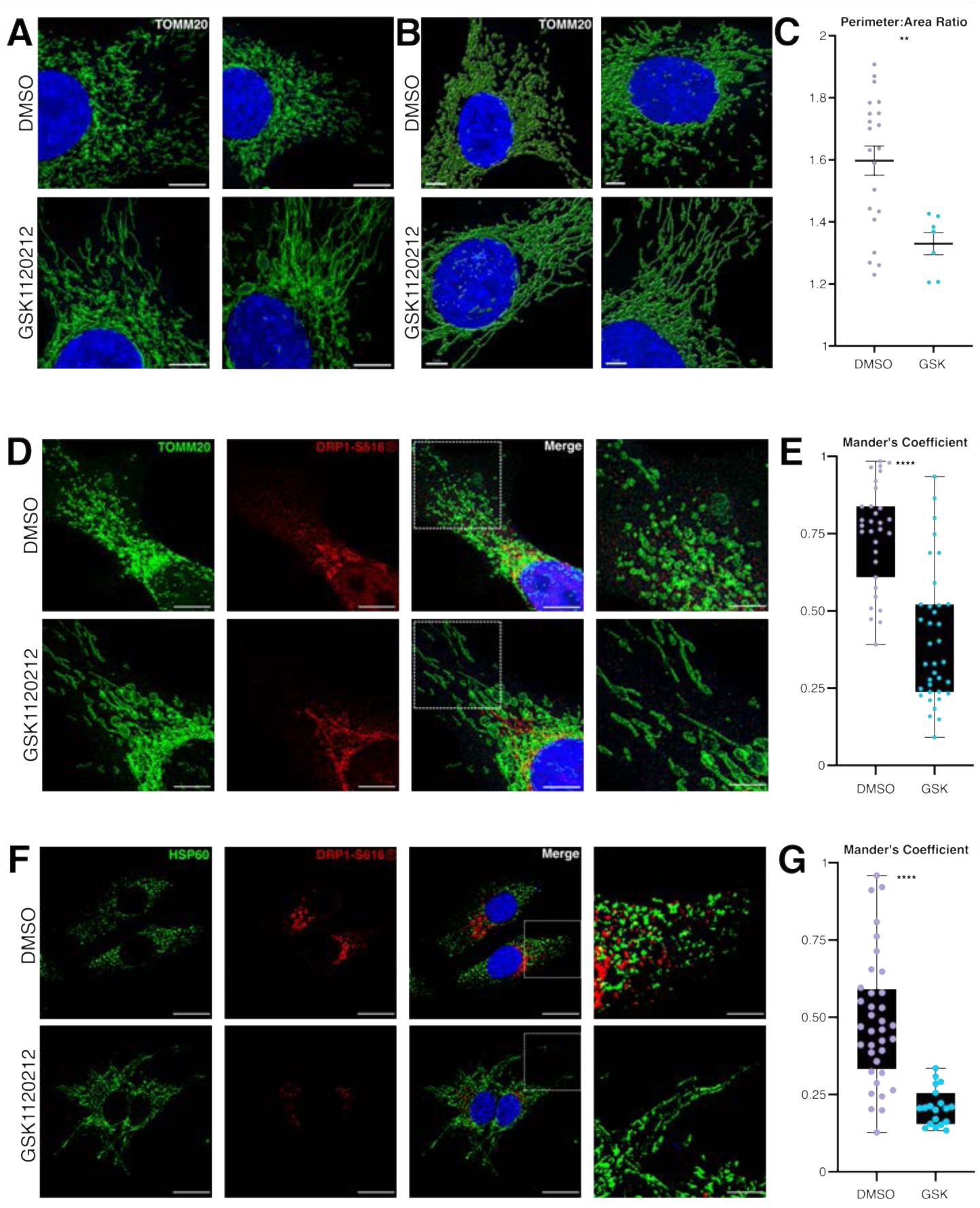
Oncogenic MAPK regulated DRP1-S616Ⓟ expression is detected by recombinant 3G11. **(A)** SKMEL28 cells were treated with GSK1120212 (50 nM) for 16 hours, fixed, and stained for TOMM20 (green; mitochondria) and Hoechst 33342 (blue; nuclei). Scale bars = 10 microns. **(B)** SKMEL28 cells were studied as in *A*. Mitochondrial networks are shown as maximum intensity projections of super-resolution Leica LIGHTNING deconvolved confocal z-stacks. Imaris surface features were used to generate 3D projections to visualize changes in mitochondrial network architecture. Scales bars = 5 microns. **(C)** Mitochondrial elongation was quantified using the Perimeter:Area Ratio. **(D–E)** SKMEL28 cells were treated with GSK1120212 (50 nM) for 16 hours, fixed, and stained for TOMM20 (green), DRP1-S616Ⓟ (red), and Hoechst 33342 (blue). Red channel intensity was quantified to assess DRP1-S616Ⓟ signal intensity using Mander’s coefficient. Scale bars = 10 microns. Zoomed network images = 50 micron scale bars. **(F–G)** A375 cells were analyzed using the same treatment, staining, imaging, and quantification approach as panels *D–E*. All experiments were performed at least twice, with statistical significance determined from a minimum of two independent experiments using a one-way ANOVA test.

### 3.4. Recombinant 3G11 Detected DRP1-S616 Phosphorylation Correlates with BRAF^V600E^ disease

To explore the potential of recombinant 3G11 for immunohistochemistry (IHC) based applications, we evaluated DRP1-S616Ⓟ detection in formalin-fixed, paraffin-embedded (FFPE) human tissue samples. Prior work in melanoma showed a link between DRP1 status and BRAF^V600E^-driven disease [11,20]. Therefore, we aimed to determine whether 3G11 supports this association in patient samples.

We analyzed de-identified FFPE tissue sections obtained from the Mount Sinai Tisch Cancer Center Biorepository Shared Resource, including normal skin, benign nevi, atypical nevi, and primary melanomas. Immunohistochemistry was performed using the recombinant 3G11 and the BRAF^V600E^ mutant specific antibodies, and staining intensity was compared across lesion types. Normal skin samples served as a negative control for both DRP1-S616Ⓟ (0 positive samples / 23 total samples) and BRAF^V600E^ (0 positive samples / 23 total samples) stainings and representative images are provided **(Figure S2)**.

In benign nevi, 18/23 DRP1-S616Ⓟ positive samples (78.26%) were BRAF^V600E^ mutant; BRAF^Wt^ and BRAF^V600E^ lesions were 41.7% (5/12) and 46.2% (21/39) DRP1-S616Ⓟ positive, respectively **(Table 1)**. Among atypical nevi with the BRAF^V600E^ mutation, 17/21 samples (80.95%) were DRP1-S616Ⓟ positive; BRAF^Wt^ and BRAF^V600E^ lesions were 50% (6/12) and 81% (17/21) DRP1-S616Ⓟ positive, respectively **(Table 2)**. Lastly, in the melanoma cohort, which included 24 BRAF^Wt^ and 57 BRAF^V600E^ tumors, we detected DRP1-S616Ⓟ in 45 samples; BRAF^Wt^ and BRAF^V600E^ lesions were 20.8% (19/24) and 70.2% (40/57) DRP1-S616Ⓟ positive, respectively **(Table 3)**. These data suggest that DRP1-S616Ⓟ may be associated with clinically dysplastic lesions and primary melanoma, implicating chronic mitochondrial division as a potential mechanistic contributor to disease.

**Table 1.**
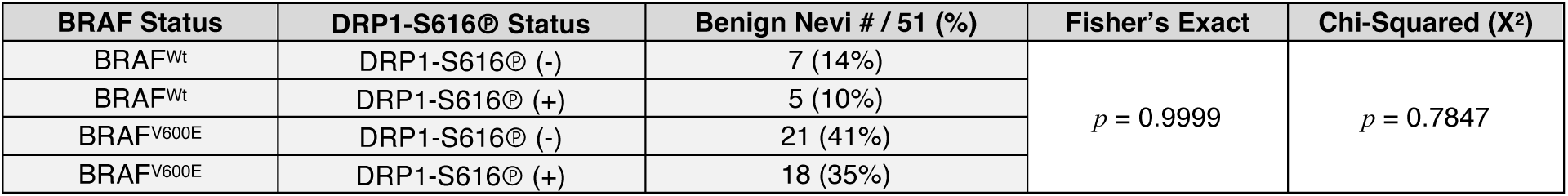
Recombinant 3G11 staining results with benign nevi sections. Immunohistochemistry was performed to detect the status of BRAF^V600E^ and DRP1-S616Ⓟ of benign nevi (51 samples). Fisher’s Exact and Chi-Squared tests determined statistical significance.

| BRAF Status | DRP1-S616® Status | Benign Nevi # / 51 (%) | Fisher’s Exact | Chi-Squared (X²) |
| --- | --- | --- | --- | --- |
| BRAF<sup>Wt</sup> | DRP1-S616® (-) | 7 (14%) | $p = 0.9999$ | $p = 0.7847$ |
| BRAF<sup>Wt</sup> | DRP1-S616® (+) | 5 (10%) |  |  |
| BRAF<sup>V600E</sup> | DRP1-S616® (-) | 21 (41%) |  |  |
| BRAF<sup>V600E</sup> | DRP1-S616® (+) | 18 (35%) |  |  |

**Table 2.** Recombinant 3G11 staining results with atypical nevi sections. Immunohistochemistry was performed to detect the status of BRAF^V600E^ and DRP1-S616Ⓟ of atypical nevi (33 samples). Fisher’s Exact and Chi-Squared tests determined statistical significance.

| BRAF Status | DRP1-S616® Status | Atypical Nevi # / 33 (%) | Fisher’s Exact | Chi-Squared (X²) |
| --- | --- | --- | --- | --- |
| BRAF<sup>Wt</sup> | DRP1-S616® (-) | 6 (18%) | $p = 0.1142$ | $p = 0.0627$ |
| BRAF<sup>Wt</sup> | DRP1-S616® (+) | 6 (18%) |  |  |
| BRAF<sup>V600E</sup> | DRP1-S616® (-) | 4 (13%) |  |  |
| BRAF<sup>V600E</sup> | DRP1-S616® (+) | 17 (51%) |  |  |

**Table 3.**
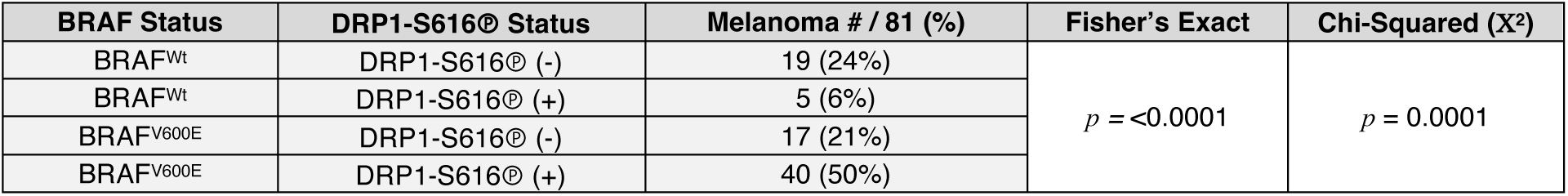
Recombinant 3G11 staining results with primary melanoma sections. Immunohistochemistry was performed to detect the status of BRAF^V600E^ and DRP1-S616Ⓟ of primary melanoma (81 samples). Fisher’s Exact and Chi-Squared tests determined statistical significance.

| BRAF Status | DRP1-S616® Status | Melanoma # / 81 (%) | Fisher's Exact | Chi-Squared (X²) |
| --- | --- | --- | --- | --- |
| BRAF<sup>WT</sup> | DRP1-S616® (-) | 19 (24%) | $p = <0.0001$ | $p = 0.0001$ |
| BRAF<sup>WT</sup> | DRP1-S616® (+) | 5 (6%) |  |  |
| BRAF<sup>V600E</sup> | DRP1-S616® (-) | 17 (21%) |  |  |
| BRAF<sup>V600E</sup> | DRP1-S616® (+) | 40 (50%) |  |  |

## 4. Discussion

In this study, we developed and characterized 3G11, a human recombinant monoclonal DRP1-S616Ⓟ antibody. We describe that 3G11 is a sensitive and reliable tool for detecting DRP1-S616Ⓟ across multiple applications. 3G11 exhibits high specificity for recombinant DRP1-S616Ⓟ protein (Figure 2), successfully detects endogenous DRP1-S616Ⓟ in cell lysates and isolated organelles via western blot (Figure 2), and effectively tracked dynamic changes in DRP1-S616Ⓟ in response to oncogenic MAPK inhibition using immunofluorescence (Figure 3). Most notably, in patient tissue sections, 3G11 also positivity revealed a correlation between DRP1-S616Ⓟ and BRAF^V600E^ benign nevi, atypical nevi, and malignant primary melanocytic lesions (Figure 4).

Recombinant 3G11 readily detects DRP1-S616Ⓟ in archival and recent FFPE tissues, immunoblots, and immunofluorescence assays. This versatility expands the tools available for studying mitochondrial dynamics in diverse cancer models and human samples. This supports explorations aimed at understanding how DRP1-S616Ⓟ influences metabolic rewiring, therapy responses, and disease progression as chronic mitochondrial division is implicated in the mechanistic control of these phenotypes [15,24–26]. DRP1-S616Ⓟ status is also linked to tumor types beyond melanoma, so broader applications of 3G11 have the potential to define shared mitochondrial vulnerabilities in multiple tumor types [12–17]. Future studies should also examine DRP1-S616Ⓟ in receptor tyrosine kinase (*e.g.,* EGFR^L858R^) and RAS (*e.g.,* N-RAS^Q61R^) mutant tumors to define a more complete biological and translational scope.

The strength of this association in Tables 1–3 reveals the impact of DRP1-S616Ⓟ as a molecular readout of mitochondrial division in BRAF^V600E^ tumors; furthermore, DRP1-S616Ⓟ is also elevated in RAS^G12V^ cancer models [11,12]. Both BRAF^V600E^ and RAS^G12V^ tumors share metabolic features that include increased glycolytic activity, eroded oxidative phosphorylation, and persistent mitochondrial fragmentation [11, 12]. These related observations highlight the utility of studying DRP1-S616Ⓟ-dependent chronic mitochondrial division for deeper insights into fundamental cancer mechanisms and prognosis.

Compared with most commercially available DRP1-S616Ⓟ antibodies commonly used in the literature, the new human-specific recombinant antibody offers several important advantages for the study of mitochondrial division. Most antibodies targeting DRP1-S616Ⓟ are polyclonal preparations generated against undefined phosphopeptides using standard hybridomas, resulting in heterogeneous antibody populations with variable affinity and epitope recognition. This heterogeneity can lead to lot-to-lot variability, reduced quantitative reliability, and increased susceptibility to cross-reactivity with closely related phospho-epitopes. In contrast, recombinant production of a defined VH/L human-sequence monoclonal DRP1-S616Ⓟ antibody ensures molecular uniformity, batch-to-batch consistency, stable long-term availability throughout research program longevity, and is more economical. The single human-specific epitope further improves performance in complex human samples, including FFPE tissues, where polyclonal reagents frequently exhibit elevated background or inconsistent staining and are not recommended. In summary, recombinant 3G11 demonstrates robust and reproducible detection of DRP1-S616Ⓟ across immunoblotting, immunofluorescence, and histologic platforms, thereby enhancing experimental rigor throughout a complex study and supporting its use for integrated mechanistic and translational investigations of mitochondrial division in cancer and related disease contexts.

## 5. Conclusions

This work introduces 3G11, a human recombinant monoclonal antibody that detects DRP1-S616Ⓟ with high sensitivity and specificity across multiple experimental platforms. 3G11 performed robustly in immunoblotting, immunofluorescence, and FFPE tissue sections. Using this reagent, we identified a consistent association between DRP1-S616Ⓟ and BRAF^V600E^ across benign, atypical, and malignant melanocytic lesions. These findings support prior reports linking oncogenic MAPK signaling to sustained DRP1-S616Ⓟ and underscore the relevance of this phosphorylation event as a molecular feature of oncogenic MAPK-driven tumors. Collectively, 3G11 represents a specific, versatile, renewable, and cost-effective tool for studying mitochondrial division, with strong potential for translational and clinical applications.

## Supplementary Materials

The following supplementary information can be downloaded at: https://www.mdpi.com/article/doi/s1, Figure S1: Dot blot screening of DRP1-S616Ⓟ anti-sera; Figure S2: Representative immunohistochemical stainings for DRP1-S616Ⓟ and BRAF^V600E^ in tissue cohorts.

## Author Contributions

Conceptualization, J.E.C., M.N.S., and Y.C.; methodology, S.T.N., Y.C., J.A.D., R.F.R., M.S.; software, S.T.N. and K.D.S.; validation, S.T.N., J.A.D., and Y.C.; formal analysis, S.T.N and K.S.; investigation, S.T.N., Y.C., J.K., A.M., J.F.H., S.G.B.; resources, M.N.S.; data curation, S.T.N. and K.S.; writing—original draft preparation, S.T.N.; writing—review, rewriting, and editing, S.T.N. and J.E.C.; visualization, S.T.N., J.D.G.; supervision, J.E.C., S.M., M.S.; project administration, S.T.N.; funding acquisition, M.S., M.N.S., and J.E.C. All authors have read and agreed to the published version of the manuscript.

## Funding

This work was supported by NIH grants: R01-CA237264 (J.E.C.), R01-CA267696 (J.E.C.), R01-CA271346 (J.E.C.), and K22-CA218480 (M.N.S.); a Collaborative Pilot Award from the Melanoma Research Alliance (J.E.C.); a Department of Defense − Congressionally Directed Medical Research Programs − Melanoma Research Program: Mid-Career Accelerator Award (ME210246; J.E.C.); an award from the National Science Foundation (2217138); a Translational Award Program from the V Foundation (T2023-010); and the Mount Sinai Tisch Cancer Cancer (P30-CA196521). The authors would like to acknowledge the Mount Sinai Tisch Cancer Center Shared Resources and the Department of Oncological Sciences for research support.

## Informed Consent Statement

Not applicable.

## Data Availability Statement

The raw data supporting the conclusions of this article will be made available by the authors on request.

## Acknowledgments

We thank the staff within the Center for Therapeutic Antibody Development at Mount Sinai for their guidance and technical support; Dr. Rachel Brody in the Mount Sinai Tisch Cancer Center Biorepository and Pathology Shared Resource for providing the skin tissue samples and for their support with the immunohistochemistry; the staff within the Mount Sinai Tisch Cancer Center Microscopy and Advanced Bioimaging Shared Resource for their guidance and technical support; Dr. Julide T. Celebi (NYU Grossman School of Medicine, New York, NY, USA) for the primary melanoma cell lines.

## Conflicts of Interest

The authors declare no conflicts of interest.

## Abbreviations

The following abbreviations are used in this manuscript:

DMSO: Dimethyl Sulfoxide
DRP1: Dynamin Related Protein 1
ERK1/2: Extracellular Regulated Kinase 1/2
IMM: Inner Mitochondrial Membrane
MAPK: Mitogen Activated Protein Kinase
MFN1/2: Mitofusin-1/2
OIS: Oncogene-Induced Senescence
OMM: Outer Mitochondrial Membrane
OPA1: Optic Atrophy 1
ELISA: Enzyme-Linked Immunosorbent Assay
VH/L: Variable Heavy and Light Chains

**Disclaimer/Publisher’s Note:** The statements, opinions and data contained in all publications are solely those of the individual author(s) and contributor(s) and not of MDPI and/or the editor(s). MDPI and/or the editor(s) disclaim responsibility for any injury to people or property resulting from any ideas, methods, instructions or products referred to in the content.

## 6. Supplementary Materials

**Figure S1. (Related to Figure 1).**
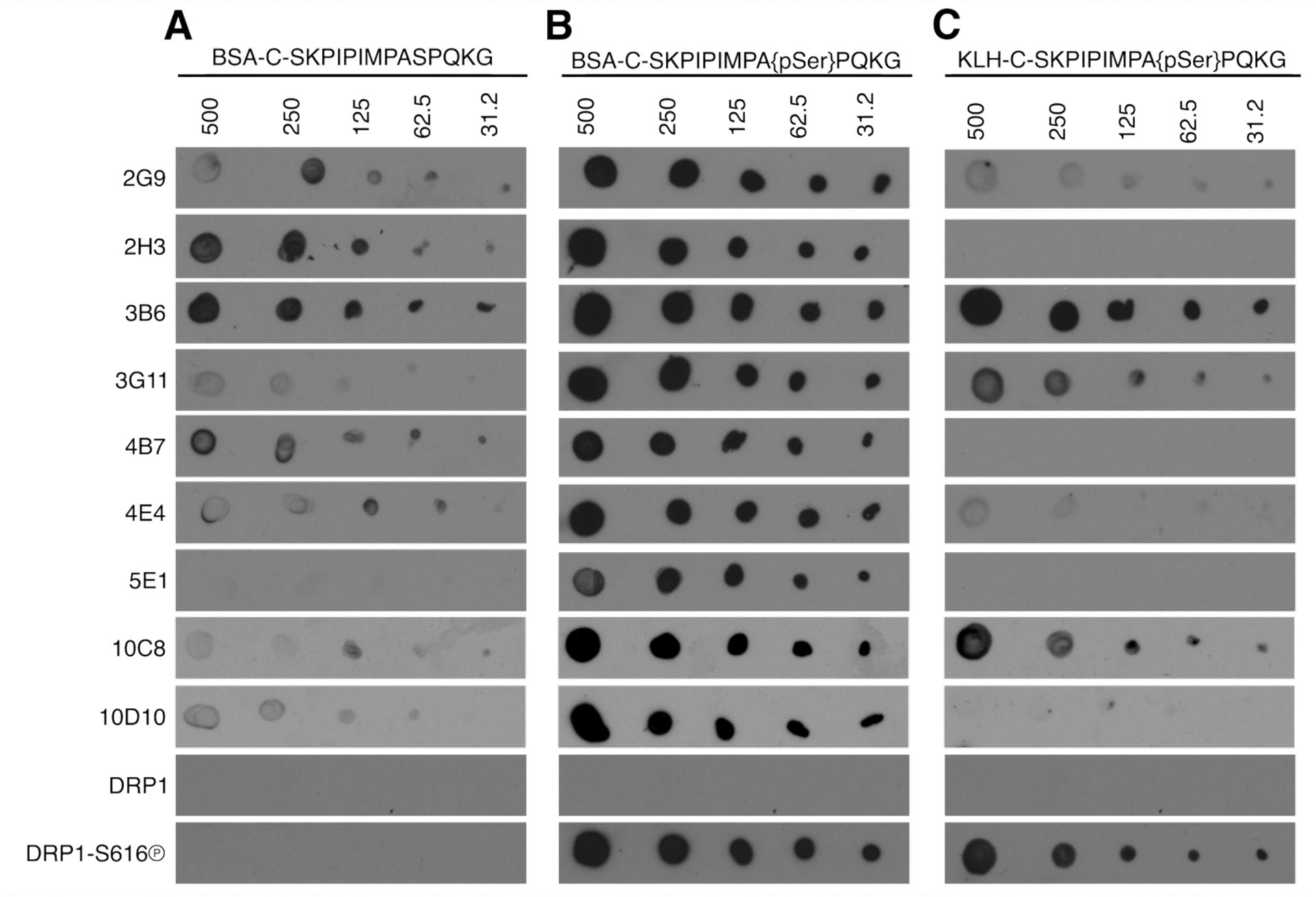
Dot blot screening of DRP1-S616Ⓟ anti-sera. **(A–C)** Dot blot analysis of anti-sera assessing specificity for the KLH-conjugated DRP1-S616Ⓟ peptide. Indicated peptides were spotted and dried onto nitrocellulose membranes at decreasing amounts (ng/dot) and detected by standard enhanced chemiluminescence. The bottom two rows of dot blots were probed using commercially available purified rabbit polyclonal DRP1 and DRP1-S616Ⓟ antibodies.

**Figure S2 (Related to Tables 1–3).**
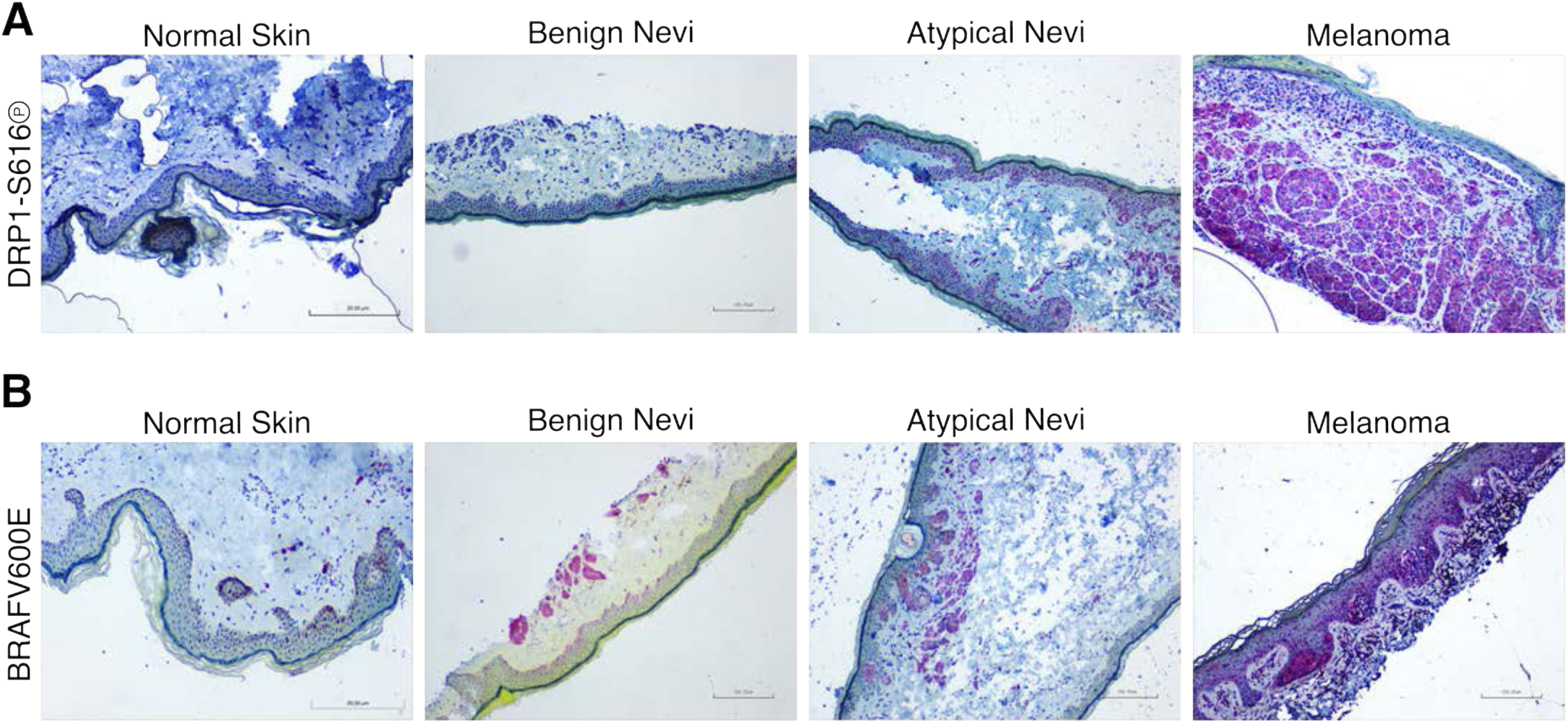
Immunohistochemical stainings for DRP1-S616Ⓟ and BRAF^V600E^ in patient cohorts. **(A)** Representative images of recombinant 3G11 staining in normal skin, benign nevi, atypical nevi, and primary melanoma lesions. Normal skin serves as the control tissue and defines negative staining. Benign nevi show limited and diffuse staining. Atypical nevi show increased staining relative to benign nevi. Primary melanoma lesions show strong and diffuse staining. **(B)** Representative images of BRAF^V600E^ staining in normal skin, benign nevi, atypical nevi, and primary melanoma lesions. Normal skin serves as the control tissue and defines negative staining. Benign nevi showed broad BRAF^V600E^ positivity patterns including no staining, diffuse, and/or strong intensity; we presume due to the heterogeneous nature of benign nevi and a combination of lesions that were stable versus a set that potentially progressed to dysplastic and/or primary disease. Atypical nevi show increased staining compared to benign nevi. Primary melanoma lesions show robust cytosolic staining.

## Notes

### Competing Interest Statement

The authors have declared no competing interest.

## References

1. Adhikary, A., Mukherjee, A., Banerjee, R., & Nagotu, S. (2023). DRP1: At the Crossroads of Dysregulated Mitochondrial Dynamics and Altered Cell Signaling in Cancer Cells. ACS Omega, 8(48), 45208–45223. 10.1021/acsomega.3c06547

2. Banerjee, R., Mukherjee, A., & Nagotu, S. (2022). Mitochondrial dynamics and its impact on human health and diseases: inside the DRP1 blackbox. J Mol Med (Berl), 100(1), 1–21. 10.1007/s00109-021-02150-7

3. Collier, J. J., Oláhová, M., McWilliams, T. G., & Taylor, R. W. (2023). Mitochondrial signalling and homeostasis: from cell biology to neurological disease. Trends in Neurosciences, 46(2), 137–152. 10.1016/j.tins.2022.12.001

4. Tezze, C., Romanello, V., Desbats, M. A., Fadini, G. P., Albiero, M., Favaro, G., Ciciliot, S., Soriano, M. E., Morbidoni, V., Cerqua, C., Loefler, S., Kern, H., Franceschi, C., Salvioli, S., Conte, M., Blaauw, B., Zampieri, S., Salviati, L., Scorrano, L., & Sandri, M. (2017). Age-Associated Loss of OPA1 in Muscle Impacts Muscle Mass, Metabolic Homeostasis, Systemic Inflammation, and Epithelial Senescence. Cell Metab, 25(6), 1374–1389.e1376. 10.1016/j.cmet.2017.04.021

5. Banerjee, R., Mukherjee, A., & Nagotu, S. (2022). Mitochondrial dynamics and its impact on human health and diseases: inside the DRP1 blackbox. J Mol Med (Berl), 100(1), 1–21. 10.1007/s00109-021-02150-7

6. Khatun, J., Gelles, J. D., & Chipuk, J. E. (2024). Dynamic death decisions: How mitochondrial dynamics shape cellular commitment to apoptosis and ferroptosis. Dev Cell, 59(19), 2549–2565. 10.1016/j.devcel.2024.09.004

7. Jin, J.-y., Wei, X.-x., Zhi, X.-l., Wang, X.-h., & Meng, D. (2021). Drp1-dependent mitochondrial fission in cardiovascular disease. Acta Pharmacologica Sinica, 42(5), 655–664. 10.1038/s41401-020-00518-y

8. Poulikakos, P. I., Sullivan, R. J., & Yaeger, R. (2022). Molecular Pathways and Mechanisms of BRAF in Cancer Therapy. Clin Cancer Res, 28(21), 4618–4628. 10.1158/1078-0432.Ccr-21-2138

9. Teixido, C., Castillo, P., Martinez-Vila, C., Arance, A., & Alos, L. (2021). Molecular Markers and Targets in Melanoma. Cells, 10(9). 10.3390/cells10092320

10. Ascierto, P. A., Kirkwood, J. M., Grob, J. J., Simeone, E., Grimaldi, A. M., Maio, M., Palmieri, G., Testori, A., Marincola, F. M., & Mozzillo, N. (2012). The role of BRAF V600 mutation in melanoma. J Transl Med, 10, 85. 10.1186/1479-5876-10-85

11. Serasinghe, M. N., Wieder, S. Y., Renault, T. T., Elkholi, R., Asciolla, J. J., Yao, J. L., Jabado, O., Hoehn, K., Kageyama, Y., Sesaki, H., & Chipuk, J. E. (2015). Mitochondrial division is requisite to RAS-induced transformation and targeted by oncogenic MAPK pathway inhibitors. Mol Cell, 57(3), 521–536. 10.1016/j.molcel.2015.01.003

12. Kashatus, J. A., Nascimento, A., Myers, L. J., Sher, A., Byrne, F. L., Hoehn, K. L., Counter, C. M., & Kashatus, D. F. (2015). Erk2 phosphorylation of Drp1 promotes mitochondrial fission and MAPK-driven tumor growth. Mol Cell, 57(3), 537–551. 10.1016/j.molcel.2015.01.002

13. Rehman, J., Zhang, H. J., Toth, P. T., Zhang, Y., Marsboom, G., Hong, Z., Salgia, R., Husain, A. N., Wietholt, C., & Archer, S. L. (2012). Inhibition of mitochondrial fission prevents cell cycle progression in lung cancer. The FASEB Journal, 26(5), 2175–2186. 10.1096/fj.11-196543

14. Inoue-Yamauchi, A., & Oda, H. (2012). Depletion of mitochondrial fission factor DRP1 causes increased apoptosis in human colon cancer cells. Biochem Biophys Res Commun, 421(1), 81–85. 10.1016/j.bbrc.2012.03.118

15. Xie, Q., Wu, Q., Horbinski, C. M., Flavahan, W. A., Yang, K., Zhou, W., Dombrowski, S. M., Huang, Z., Fang, X., Shi, Y., Ferguson, A. N., Kashatus, D. F., Bao, S., & Rich, J. N. (2015). Mitochondrial control by DRP1 in brain tumor initiating cells. Nature Neuroscience, 18(4), 501–510. 10.1038/nn.3960

16. Huang, T. L., Chang, C. R., Chien, C. Y., Huang, G. K., Chen, Y. F., Su, L. J., Tsai, H. T., Lin, Y. S., Fang, F. M., & Chen, C. H. (2022). DRP1 contributes to head and neck cancer progression and induces glycolysis through modulated FOXM1/MMP12 axis. Mol Oncol, 16(13), 2585–2606. 10.1002/1878-0261.13212

17. Zhao, J., Zhang, J., Yu, M., Xie, Y., Huang, Y., Wolff, D. W., Abel, P. W., & Tu, Y. (2013). Mitochondrial dynamics regulates migration and invasion of breast cancer cells. Oncogene, 32(40), 4814–4824. 10.1038/onc.2012.494

18. Siegel, R. L., Kratzer, T. B., Giaquinto, A. N., Sung, H., & Jemal, A. (2025). Cancer statistics, 2025. CA: A Cancer Journal for Clinicians, 75(1), 10–45. 10.3322/caac.21871

19. García-Gómez, R., Bustelo, X. R., & Crespo, P. (2018). Protein-Protein Interactions: Emerging Oncotargets in the RAS-ERK Pathway. Trends Cancer, 4(9), 616–633. 10.1016/j.trecan.2018.07.002

20. Wieder, S. Y., Serasinghe, M. N., Sung, J. C., Choi, D. C., Birge, M. B., Yao, J. L., Bernstein, E., Celebi, J. T., & Chipuk, J. E. (2015). Activation of the Mitochondrial Fragmentation Protein DRP1 Correlates with BRAF(V600E) Melanoma. J Invest Dermatol, 135(10), 2544–2547. 10.1038/jid.2015.196

21. Renault, T. T., Floros, K. V., Elkholi, R., Corrigan, K. A., Kushnareva, Y., Wieder, S. Y., Lindtner, C., Serasinghe, M. N., Asciolla, J. J., Buettner, C., Newmeyer, D. D., & Chipuk, J. E. (2015). Mitochondrial shape governs BAX-induced membrane permeabilization and apoptosis. Mol Cell, 57(1), 69–82. 10.1016/j.molcel.2014.10.028

22. Rochon, K., Bauer, B. L., Roethler, N. A., Buckley, Y., Su, C.-C., Huang, W., Ramachandran, R., Stoll, M. S. K., Yu, E. W., Taylor, D. J., & Mears, J. A. (2024). Structural basis for regulated assembly of the mitochondrial fission GTPase Drp1. Nature Communications, 15(1), 1328. 10.1038/s41467-024-45524-4

23. Singh, S., & Sharma, S. (2017). Dynamin-related protein-1 as potential therapeutic target in various diseases. Inflammopharmacology, 25(4), 383–392. 10.1007/s10787-017-0347-y Teixido, C., Castillo, P., Martinez-Vila, C., Arance, A., & Alos, L. (2021). Molecular Markers and Targets in Melanoma. Cells, 10(9). 10.3390/cells10092320

24. Sharmin, S., Kashatus, J. A., Adair, S. J., Bauer, T. W., & Kashatus, D. F. (2025). Abstract LB283: Reactivation of Drp1S616 phosphorylation by c-MYC-CDK4/6 signaling axis plays a functional role in resistance to MEK inhibition in pancreatic cancer cells. Cancer Research, 85(8_Supplement_2), LB283–LB283. 10.1158/1538-7445.Am2025-lb283

25. Sami Alkafaas, S., Obeid, O. K., Ali Radwan, M., Elsalahaty, M. I., Samy ElKafas, S., Hafez, W., Janković, N., & Hessien, M. (2024). Novel insight into mitochondrial dynamin-related protein-1 as a new chemo-sensitizing target in resistant cancer cells. Bioorganic Chemistry, 150, 107574. 10.1016/j.bioorg.2024.107574

26. Serasinghe, M. N., Gelles, J. D., Li, K., Zhao, L., Abbate, F., Syku, M., Mohammed, J. N., Badal, B., Rangel, C. A., Hoehn, K. L., Celebi, J. T., & Chipuk, J. E. (2018). Dual suppression of inner and outer mitochondrial membrane functions augments apoptotic responses to oncogenic MAPK inhibition. Cell Death Dis, 9(2), 29. 10.1038/s41419-017-0044-1

